# Intratumoral injection of melatonin enhances tumor regression in cell line-derived and patient-derived xenografts of head and neck cancer by increasing mitochondrial oxidative stress

**DOI:** 10.1101/2023.07.17.549311

**Authors:** Laura Martinez-Ruiz, Javier Florido, César Rodriguez-Santana, Alba López-Rodríguez, Ana Guerra-Librero, Beatriz I. Fernández-Gil, Patricia García-Tárraga, José Manuel Garcia-Verdugo, Felix Oppel, Holger Sudhoff, David Sánchez-Porras, Amadeo Ten-Steve, José Fernández-Martínez, Pilar González-García, Iryna Rusanova, Darío Acuña-Castroviejo, Víctor S. Carriel, Germaine Escames

**Affiliations:** Institute of Biotechnology, Biomedical Research Center, Health Sciences Technology Park, University of Granada, Granada, Spain; Department of Physiology, Faculty of Medicine, University of Granada, Granada, Spain; Centro de Investigación Biomédica en Red Fragilidad y Envejecimiento Saludable (CIBERFES), Instituto de Investigación Biosanitaria (Ibs), Granada, San Cecilio University Hospital, Granada, Spain; Department of Neurologic Surgery, Mayo Clinic, Jacksonville, USA; Department of Histology, Tissue Engineering Group, Faculty of Medicine, University of Granada, Granada, Spain; Instituto de Investigación Biosanitaria ibs. GRANADA, Granada, Spain; Department of Biochemistry and Molecular Biology I, Faculty of Science, University of Granada, Granada, Spain; Department of Otolaryngology, Head and Neck Surgery, University Hospital OWL of Bielefeld University, Campus Klinikum Bielefeld Mitte, Teutoburger Str. 50, 33604, Bielefeld, Germany; Cavanilles Institute of Biodiversity and Evolutionary Biology, University of Valencia, Valencia, Spain; Biomedical Imaging Research Group (GIBI230-PREBI), La Fe Health Research Institute and Imaging La Fe node at Distributed Network for Biomedical Imaging, Unique Scientific and Technical Infrastructures, Valencia, Spain

**Keywords:** Melatonin, intratumoral injection, patient-derived xenograft, head and neck cancer, mitochondria, reactive oxygen species

## Abstract

Head and neck squamous cell carcinoma present a high mortality rate. Melatonin has been shown to have oncostatic effects in different types of cancers. However, inconsistent results have been reported for *in vivo* applications. Consequently, an alternative administration route is needed to improve bioavailability and establish the optimal dosage of melatonin for cancer treatment. On the other hand, the use of patient-derived tumor models has transformed the field of drug research because they reflect the heterogeneity of patient tumor tissues. In the present study, we explore mechanisms for increasing melatonin bioavailability in tumors and investigate its potential as an adjuvant to improve the therapeutic efficacy of cisplatin in the setting of both xenotransplanted cell lines and primary human HNSCC. We analyzed the effect of two different formulations of melatonin administered subcutaneously or intratumorally in Cal-27 and SCC-9 xenografts and in patient-derived xenografts. Melatonin effects on tumor mitochondrial metabolism was also evaluated as well as melatonin actions on tumor cell migration. In contrast to the results obtained with the subcutaneous melatonin, intratumoral injection of melatonin drastically inhibited tumor progression in HNSCC-derived xenografts, as well as in patient-derived xenografts. Interestingly, intratumoral injection of melatonin potentiated CDDP effects, decreasing Cal-27 tumor growth. We demonstrated that melatonin increases ROS production and apoptosis in tumors, targeting mitochondria. Melatonin also reduces migration capacities and metastasis markers. These results illustrate the great clinical potential of intratumoral melatonin treatment and encourage a future clinical trial in cancer patients to establish a proper clinical melatonin treatment.

## 1. Introduction

Head and neck squamous cell carcinoma (HNSCC) is one of the most common malignancies worldwide. Due to its heterogeneous biology, treatments are complex, and most patients often require multimodal intervention, such as surgery, radiotherapy, and/or chemoradiotherapy [1]. However, the prognosis of HNSCC remains poor, with a high mortality rate [2,3]. There is a high incidence of resistance to radio and chemotherapy, which contributes to tumor progression and metastasis [4,5]. Therefore, effective strategies to counteract resistance and improve clinical outcome are urgently needed [6].

Melatonin (N-acetyl-5-methoxytryptamine) has been shown to have oncostatic effects in different types of cancers [4,7,8]. Previous studies have reported that melatonin regulates mitochondrial function [9] and drives apoptosis by increasing mitochondrial reactive oxygen species (ROS) in HNSCC [9–14]. However, despite multiple studies demonstrating the *in vitro* efficacy of melatonin, some studies have shown that the use of melatonin alone exerts a lesser or no effect on tumor proliferation *in vivo*, especially in HNSCC [12,15]. Moreover, the only clinical trial focusing on the effects of melatonin in HNSCC was published without any significant results [16], possibly due to the low bioavailability of melatonin in the tumor [15]. Therefore, it is necessary to search for an alternative administration route to improve bioavailability and establish the optimal dosage for cancer treatment [15].

Moreover, the low effectiveness of many anticancer drugs investigated *in vivo* is a result of inadequate use of existing preclinical models [17]. Established cancer cell lines have traditionally been the most used models to study tumor biology and pharmacogenomics. Nevertheless, these models have important limitations, as they only represent a subpopulation of the original tumor and are largely homogenous [18]. Interestingly, patient-derived tumor models reflect the heterogeneity of real patient tumor tissues [19] and encompass multiple factors, such as histological structures, malignant genotypes, and phenotypes [20]. Therefore, patient-derived tumor xenografts (PDXs) are a better alternative for studying cancer biology and drug sensitivity [18]. Nevertheless, the number of studies investigating the effect of melatonin on xenotransplanted primary human cancer cells has been limited [21,22].

In the present study, we explore mechanisms for increasing melatonin bioavailability in tumors and its potential as an adjuvant to improve the therapeutic efficacy of cisplatin (CDDP) in the setting of both xenotransplanted cell lines and primary human HNSCC. Interestingly, our study reveals an unexpected role of high concentrations of melatonin, which results in complete regression of the tumor and decreased metastasis. Therefore, melatonin may serve as a potential therapy for HNSCC.

## 2. Materials and methods

### 2.1. Cell culture and treatment

Two standard HNSCC cell lines were used in this study: Cal-27 (ATCC: CRL-2095) and SCC-9 (ATCC: CRL-1629). Both cell lines were obtained from the Cell Bank of the Scientific Instrumentation Centre at University of Granada. A patient-derived cell culture (PDC) was also used. This primary cell culture was obtained from a human biopsy of head and neck cancer tissue in Dr. Oppel’s laboratory, where it was previously characterized [23]. Cells were treated with melatonin at 500 µM or 1000 µM. Vehicle was added to the control group. Melatonin (33457-24, Fagron Ibérica S.A.U., Terrassa, Spain) stock solution was prepared with 15% propane-1,2-diol (PG; 24414.296, VWR) in Dulbecco’s phosphate-buffered saline (DPBS; 14190094, Life Technologies, Carlsbad, California, USA).

### 2.2. Animal models

This study was conducted using 5-to 6-week-old athymic (nu/nu) nude mice (Janvier Labs, France) for established cancer cell line xenografting and NOD scid gamma (NSG) mice for PDXs. The animals were maintained in the University’s facility in a 12:12-hr light/ dark cycle (lights on at 07:00 hr), and at 22°C and on regular chow and tap water. All mice were housed in appropriate sterile filter-capped cages and fed and given water ad libitum.

All experiments were performed following a protocol approved by the Institutional Animal Care and Use Committee of the University of Granada (procedures 15/10/2020/119), developed in accordance with the European Convention for the Protection of Vertebrate Animals used for Experimental and Other Scientific Purposes (CETS # 123) and Spanish law (R.D. 53/2013). Primary tumors were obtained from medically indicated surgeries conducted at Klinikum Bielefeld with informed consent of the patients, according to the declaration of Helsinki and as approved by the ethics committee of the Ruhr-University Bochum (AZ 2018– 397-1).

For standard cancer cell line xenografts, two HNSCC cell lines were used: Cal-27 and SCC-9. Cell viability was assessed using the trypan blue exclusion test. The cell suspension was subcutaneously transplanted into the left flanks of the mice at a 4 × 10^6^/0.2 mL concentration in DMEM (31053-044, ThermoFisher Scientific) and Matrigel Basement Membrane Matrix (1:1) (354234, VWR).

For PDX, tumors were obtained from medically indicated surgeries conducted at Klinikum Bielefeld with the informed consent of the patients according to the Declaration of Helsinki and as approved by the ethics committee of Ruhr-University Bochum (AZ 2018– 397-1). Tumor pieces were digested using 5–10 mL of a 2.5 mg/mL Collagenase NB4 standard (S1745401 Nordmark Biochemicals, Uetersen, Germany) solution in PBS + 3 mM CaCl_2_ for 2 hours at 37°C. After digestion, purified cells/cell aggregates were washed in PBS and frozen in 5% dimethyl sulphoxide (DMSO; 23486.297, VWR, Radnor, PA, USA) in FBS until xenografting.

Mouse body weights were measured using a digital balance, and tumor sizes were determined using a vernier caliper. Tumor volume was calculated as (width × length^2^)/2.

### 2.3. Animal treatments

After xenografting, once the tumor volume reached 100–200 mm^3^, mice were randomly divided into different groups based on the treatment to be assessed. The mice were treated with and without melatonin (aMT), cisplatin (CDDP), irradiation (IR), CDDP plus melatonin (CDDP +aMT), or IR plus melatonin (IR +aMT).

Melatonin was dissolved in PG and diluted at 37.5% in saline solution. CDDP was dissolved in saline solution. Both melatonin and CDDP were filtered using a 0.2 µm pore filter (#PN 4612, Pall Corporation LifeSciences, California, USA) before administration into the mice.

Melatonin was administered subcutaneously at 300 mg/kg day (each 24 hours for 21 days) or intratumorally at 3% (w/v) (1,8 mg/100 mm^3^) each 24 hours until a maximum of 63 days. Control groups received vehicle. The intratumoral melatonin injectable is under patent (P201730598).

CDDP was injected intraperitoneally at 4 mg/kg day once per week. In the group receiving CDDP plus aMT, melatonin was administered at 300 mg/kg day subcutaneously (each 24 hours for 21 days) or intratumorally at 3% (each 24 hours for 49 days).

Mice were irradiated at 4 Gy in a steel cylinder with an appropriate hole for tumors after anesthetizing them via intraperitoneal injection of equitesin. The irradiation was emitted with an X-ray tube (YXLON, model Y, Tu 320-D03) using a voltage of 239 kV, working current of 13 mA, focus of irradiation 5 mm in diameter, 0.32 mm Cu filter system, target distance of 10 cm, irradiation field of 78.54 mm^2^, and delivered dosage of 1.502 ± 0.05% Gy/min. In the group of melatonin plus IR, melatonin was administered subcutaneously at 300 mg/kg every 24 hours for 7 days, and the first dose of melatonin was administered 1 day before IR.

### 2.4. Magnetic resonance imaging

Mice were anesthetized with isoflurane vaporized with O_2_. The isoflurane concentration was set at 3.0% for induction and at 1.0–2.0% for maintenance. The animal was in a prone position for transverse acquisition covering the whole volume of the tumors. This planning was made from the sagittal and coronal scout. The acquisition protocol consisted of a T2-weighted turbo spin-echo multi-slice sequence (T2W MS TSE) with a field of view (FOV) of 50×50×19 mm on the axial plane (27 slices), voxel size 0.15×0.15×0.7 mm, TE of 97 ms, and TR of 2800. All images were obtained in Philips Achieva 3.0 TX (Amsterdam, the Netherlands) using an 8 Channel Sense Wrist Coil.

### 2.5. Histology

For histological analyses, animals were subjected to intracardiac perfusion with 3.7– 4% neutral buffered paraformaldehyde (PFA). Once perfused, tumors were fixed in PFA for 24 hours and then longitudinally sectioned for histological processing and analysis. Selected samples were dehydrated in ethanol, cleared in xylol, and embedded in paraffin. Histological sections of 5 µm were dewaxed, hydrated, and stained according to the protocol.

Hematoxylin-eosin staining was used for morphological evaluation of tumor progression. The collagen network was assessed by the picrosirius (PS) histochemical method, proteoglycans and mucopolysaccharides (produced by adenosquamous cell clusters) by Alcian Blue (AB; pH 2.5) [12,24], proliferating neoplastic cells using antibodies against Ki-67 protein (MAD-000310QD, Vitro SA, Madrid, Spain), and apoptosis by TUNEL assay (G3250, Promega, Madison, WI, USA) following the manufacturers’ instructions. Images were analyzed by ImageJ software.

To quantitatively determine the tumor encapsulation, the tumor/capsule ratio was calculated using PS-stained sections and Image J Software (Free Software Foundation Inc., Boston, MA, USA). In this way, the whole tumor area, including the PS-positive capsule, was determined and the percentage of area occupied by the capsule obtained. The same quantitative approach was applied to determine the percentage of AB-positive areas occupied by mucopolysaccharides. These quantitative approaches were conducted following previously described procedures. In addition, the cell proliferation index was calculated in Ki-67-stained sections as described previously [25]. For TUNEL assay, after perfusion of the mice, tumors were carefully fixed in 4% PFA, dehydrated in 30% sucrose, and embedded in optimal cutting temperature (OCT, 331603E, VWR). Hoechst (33342, Invitrogen, ThermoFisher Scientific) was used to stain the nuclei. Images were acquired with a fluorescence microscope (Nikon Eclipse Ni-U microscope). Quantification of TUNEL+ per field was achieved using five random fields per slide.

### 2.6. Melatonin determination by HPLC

Melatonin was extracted with trichloromethane using the SPD 2010 Speed Vac System (ThermoFisher Scientific). The samples were then analyzed by HPLC (Shimadzu Europe GmbH, Duisburg, Germany) using a Waters Sunfire C18 column (150 × 4.5 mm, 5 µm). Melatonin fluorescence was measured using a Shimadzu RF-10A XL fluorescence detector (Shimadzu Europe GmbH) with 285-nm excitation and 345-nm emission wavelengths [26].

### 2.7. Western blot analysis

Protein extraction and Western blot analyses were performed as described previously [11]. Denatured protein samples (40 µg/fraction) were separated by sodium dodecyl sulfate polyacrylamide gel electrophoresis (SDS-PAGE) using 15% acrylamide/bis-acrylamide gels. Proteins were then wet transferred to a polyvinylidene difluoride (PVDF) membrane (IPVH00010, Sigma Aldrich). The membrane was incubated in blocking buffer consisting of 5% milk in PBS plus 0.1% Tween-20. Primary antibodies (Table S1) were diluted in blocking buffer and incubated overnight at 4°C. HRP-conjugated secondary antibodies were used according to the manufacturer’s instructions. Immunoreactions were detected using the ECL^TM^ Prime Western Blotting Detection Reagent (GE Healthcare Life Sciences, Barcelona, Spain), analyzed with the ChemiDoc MP Imaging System (BioRad), and quantified using ImageJ software. Protein band intensities were normalized to GAPDH or β-actin.

### 2.8. Measurement of LPO levels

The Bioxytech LPO-568 assay kit (OxisResearch, Portland, OR, USA) was used to assess lipid peroxidation (LPO). This kit uses malondialdehyde (MDA) and 4-hydroxy-2-nonenal (4-HNE) as markers of lipid peroxidation. After 40 minutes of incubation at 45°C, 1-methyl-2-phenylindole reacts with MAD and 4-HNE, yielding a stable chromophore with intense maximal absorbance at 586 nm.

### 2.9. Electron microscopy tumor analysis

For electron microscopy, animals were subjected to intracardiac perfusion with 2% PFA-2% glutaraldehyde (A17876, ThermoFisher Scientific) in 0.1 M phosphate buffer (PB; pH 7.4). Samples were then cut into 200-mm sections, washed with 0.1 M PB, and kept in 0.1 M PB-0.05% sodium azide at 4°C until subsequent use. Sections were post-fixed in 2% osmium tetroxide, dehydrated, and embedded in Durcupan resin (44611-44614, Sigma Aldrich). A diamond knife was used to cut semi-thin sections (1.5 mm), which were stained with 1% toluidine blue for light microscopy. Ultra-thin sections (70–80 nm) were then cut, stained with lead citrate [27] and evaluated under a FEI Tecnai G2 Spirit transmission electron microscope (FEI Europe) using a digital camera (Morada Soft Imaging System; Olympus) [28].

### 2.10. Evaluation of proliferation rate

For the analysis of proliferation rate, a single cell was seeded in each well of P96 plates. Cell division and growth were monitored by obtaining images on days 0, 5, 14, and 21 with an Olympus CKX41 Microscope (Olympus Deutschland GmbH, Hamburg, Germany). The diameters of cell aggregates were later analyzed with ImageJ software.

### 2.11. Measurement of ROS production

ROS production was measured using 100 µM 2’-7’-dichlorofluerescein diacetate (11) (DCFH-DA, D6883; Sigma-Aldrich), which was transformed within the cell to fluorescent 2’,7’-dichlorofluorescein. ROS levels were then measured with a microplate fluorescence reader FLx800 (Bio-Tek Instruments, Inc., Winooski, VT, USA) every 5 minutes for 45 minutes at 485 nm excitation and 530 nm emission.

### 2.12. Wound healing assay

Cells were seeded and treated with vehicle or melatonin at 500 or 1000 µM in a six-well plate (734-2323, VWR). Next, linear scratches were made by scraping the cell monolayer with a 10 µL pipette tip. Subsequently, images of the scratch wound were observed under an inverted phase-contrast microscope and recorded at 0, 24, and 48 hours using a camera (OlympusBX61, Tokyo, Japan). ImageJ was utilized to estimate the width of the wound. The migration rate was estimated by the percentage of scratch wound coverage, which was calculated as follows:

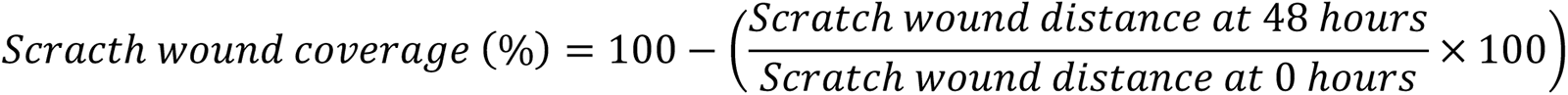

### 2.13. Transwell assay

Briefly, after melatonin treatment, while 40,000 cells were seeded in DMEM containing 0.5% FBS in the upper chambers of Transwell plates (3422, Corning, Kennebunk, USA), and the lower chambers were filled with DMEM medium containing 2% FBS. After 24 hours incubation at 37°C in a 5% CO_2_ atmosphere, migrated cells were fixed with 4% PFA, stained with DAPI (sc-24941, Santa Cruz), and counted in nine fields under a microscope (Nikon Eclipse Ni-U).

### 2.14. Measurement of nitrite levels

Nitrite levels were assessed by Griess reaction as described previously [29]. Homogenized samples were incubated with sulfanilamide and N-(1-napthyl) ethylenediamine for 20 minutes. A chromophoric azo product that absorbs strongly at 540 nm is formed by this process; absorbance was measured by a FLx800 microplate fluorescence reader.

### 2.15. Biochemical analyses

Blood samples were collected directly from the mouse heart before sacrifice. Commercial kits were used following the manufacturer’s indications. The evaluation was performed in a BS-200 Chemistry Analyzer (WN-9B100964). Commercial kits were used following the manufacturer’s instructions to analyze triglyceride (41030, Spinreact, Girona, Spain), cholesterol (41020, Spinreact), uric acid (41000 Spinreact), creatinine (1001110, Spinreact), glutamate oxaloacetate (GOT; 1001160, Spinreact), glutamate pyruvate transaminase (GPT; 1001170, Spinreact), gamma-glutamyl transferase (γ-GT; 1001185, Spinreact). and bilirubin (1001042, Spinreact) levels in plasma obtained from blood samples.

### 2.16. Data and statistical analyses

All statistical analyses were performed using Prism 8 scientific software (GraphPad Software Inc., La Jolla, CA, USA). An unpaired Student’s t-test was used to compare the differences between all groups. Data were expressed as the mean ± S.E.M. of a minimum of three independent experiments. A *p* value < .05 was considered significant.

## 3. Results

### 3.1. Subcutaneous administration of melatonin promotes tumor capsule formation but does not decrease tumor growth

Cal-27 xenograft mice were treated subcutaneously with melatonin at 300 mg/kg or vehicle daily. Surprisingly, melatonin did not result in any significant reduction in tumor volume compared to control (Fig. 1B), and these results were confirmed by MRI (Fig. 1A). These results suggest that melatonin did not reach the tumor at a sufficient concentration [12]. As suspected, melatonin-treated tumors showed a very low melatonin concentration (Fig. 1C).

Melatonin did not significantly potentiate the effects of CDDP (Fig. 1D). However, melatonin combined with IR led to greater inhibition of tumor growth than when combined with CDDP (Fig. 1F). The intratumoral melatonin concentration was very low in CDDP groups (Fig. 1E) and slightly increased in tumors treated with IR (Fig. 1G) due to the aggressiveness of IR in facilitating drug entry into the tumors [30,31].

**Fig. 1.**
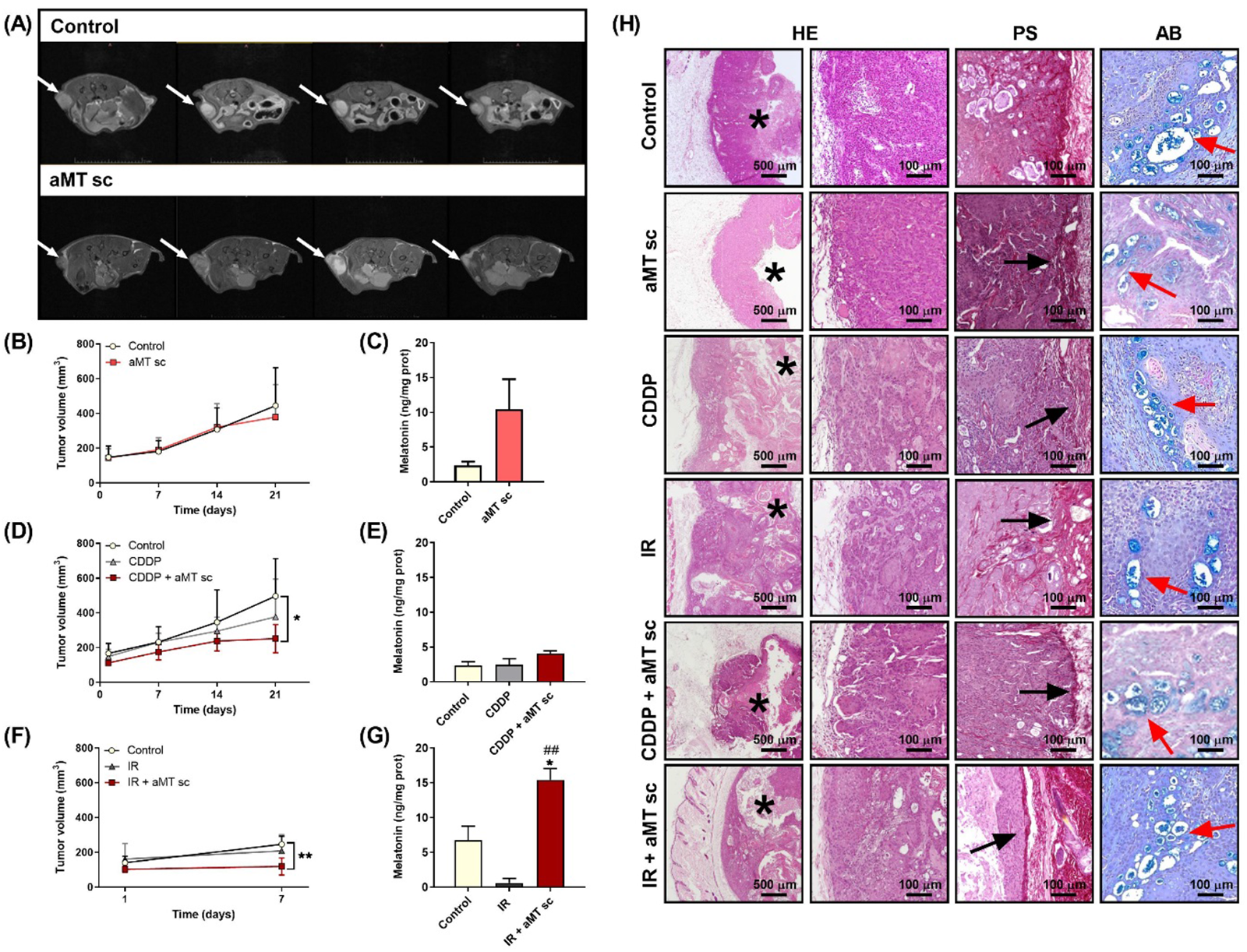
Subcutaneous administration of melatonin does not affect tumor growth in Cal-27 xenografts. (A) Representative axial T2-weigthed images obtained by MRI after 21 days of treatment in mice bearing Cal-27 tumors treated with vehicle (control) or melatonin subcutaneously (aMT sc). White arrows indicate tumor mass. (B, D, F) Tumor growth curves and (C, D, F) melatonin levels in tumors at the end of the treatments. (B, C) Treatment groups include control (mice treated with vehicle solution) and aMT sc (mice treated with melatonin at 300 mg/kg day subcutaneously). (D, E) Experimental groups comprise control, mice treated with CDDP at 4 mg/kg day once per week (CDDP), and mice treated with CDDP plus melatonin (CDDP + aMT sc). (F, G) Treatment groups involve control, mice receiving one dose of IR at 4 Gy (IR), and mice treated with the combination of IR and melatonin (IR + aMT sc). The data are presented as the means ± standard error of the mean (n=3-6 for each group). One-tailed unpaired *t*-test: \**P* < .05, \*\**P* < .01, \*\*\**P* < .001 vs. control group and ^##^ *P* < .01 vs. IR. (H) HE, PS, and AB-stained tumors from the control group and from mice treated with melatonin subcutaneously, CDDP, IR, or the combined treatments of CDDP or IR with melatonin. Images show an increase in the PS-stained collagen capsule in the groups treated with melatonin subcutaneously. Scale bar = 100 or 500 µm. Asterisks mark central tumor cysts, black arrows indicate the capsule stained red by PS, and red arrows indicate the glandular structures stained blue by AB.

Histology confirmed the development of an epidermoid carcinoma with AB-positive adenosquamous differentiation clusters (Fig. 1H). Melatonin induced a well-defined PS-positive connective capsule, but it did not reduce the tumor activity (Fig. 1H). IR or CDDP alone did not produce any significant effect compared to the control, whereas combined treatment with melatonin confirmed the impact of this molecule on encapsulation (Fig. 1H).

### 3.2. Intratumoral melatonin injection drastically inhibits tumor growth in HNSCC xenografts

To overcome the limitation of subcutaneous melatonin administration and increase melatonin levels in the tumors, intratumoral injection of melatonin was evaluated. In Cal-27 xenografts, the control group exhibited significant tumor growth from day 2 to day 41 with respect to the initial tumor volume (Fig. S1A). The mice with directly injected tumors responded markedly to the treatment with melatonin 3%, but without any significant effect with melatonin at 0.3% (Fig. 2A). Melatonin 3% exhibited a decrease in tumor volume from day 34, which was significant on day 48 (Figs. 2A and S1A), and with practically no traces of tumor after 63 days (Fig. 2A). This reduction of tumor volume correlates with a significant increase of melatonin levels in the tumors (Fig. 2B). In contrast, control tumors and tumors treated with melatonin 0.3% exceeded 1000 mm^3^ after 41 days, and the mice were sacrificed. These results were corroborated by MRI (Fig. 2C). Therefore, treatment with melatonin 3% resulted in 60%–100% regression of the tumor (Fig. 2A, C).

Results in SCC-9 xenografts were similar to those obtained in Cal-27-derived tumors. Melatonin at 3% decreased tumor growth, whereas melatonin at 0.3% had no effect (Fig. 2K). Again, this was corroborated by MRI (Fig. 2L).

In contrast to the results obtained with subcutaneous melatonin, the histology of tumors treated with intratumoral melatonin showed a decrease in tumor active areas in both cell line xenografts, which was more evident after 63 days (Figs. 2D, M). In addition, as observed previously, the formation of a collagen-rich capsule was noted in melatonin-treated tumors after 35 days, and this increased after 63 days of treatment (Figs 2D, E, N, Q). The formation of a capsule was a good signal because, as noted before, it could make the dissemination of tumor cells difficult [32,33].

Furthermore, intratumoral melatonin resulted in a significant reduction in the adenosquamous differentiation areas in Cal-27 tumors (Fig. 2D, F, G), which was not observed in subcutaneous administration (Fig. 1H), which has been associated with a better cancer prognosis [34,35].

Intratumoral melatonin decreased Ki-67, which revealed a significant reduction in cell proliferation in the active areas of treated tumors in a time-dependent manner (Figs. 2D, G, J, M, P, S). Interestingly, after melatonin treatment, the active areas remained at subcapsular tumor zones (Fig. 2D, L).

All of these findings demonstrated that the oncostatic effect of melatonin depended on reaching a high concentration in the tumors [12].

**Fig. 2.**
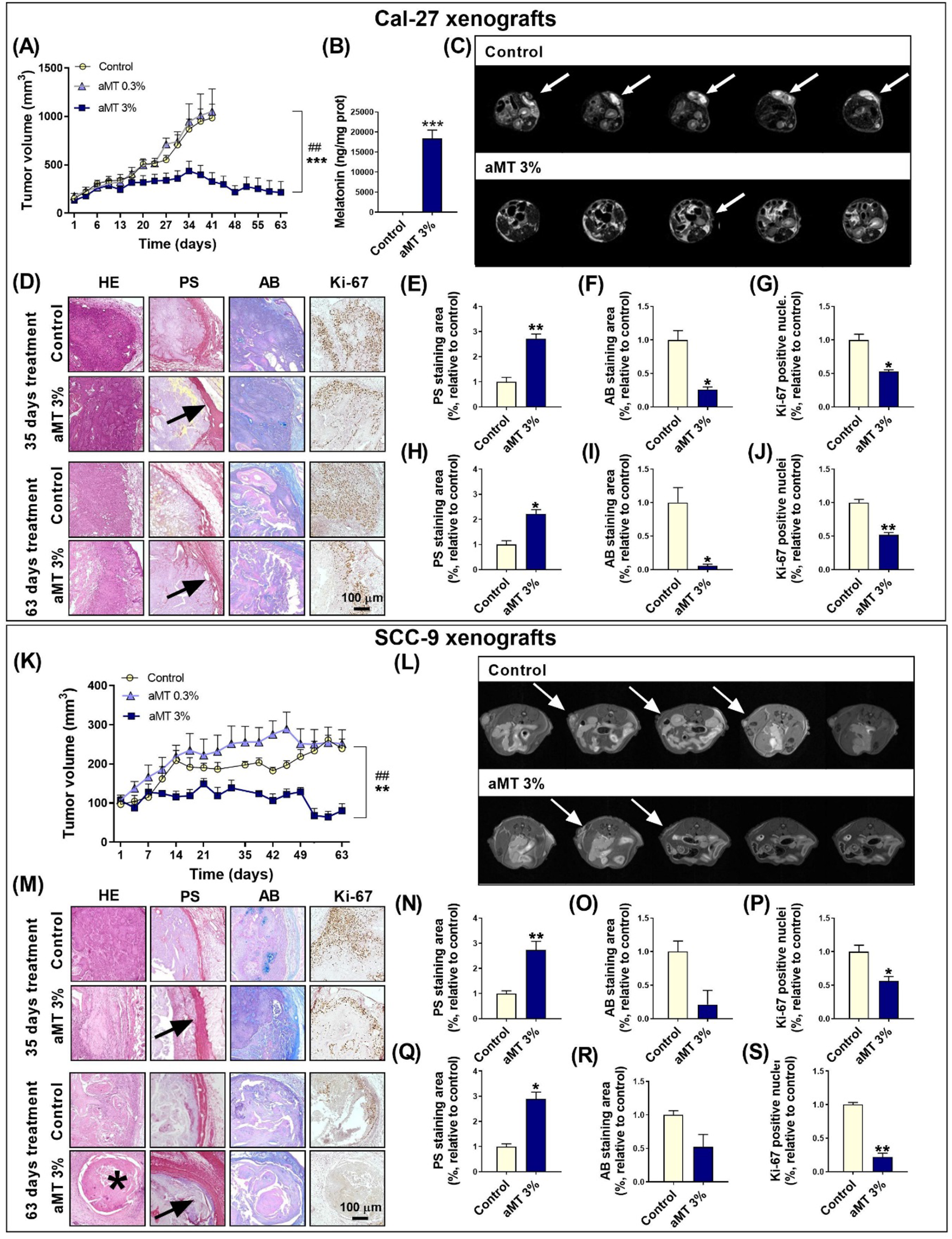
Intratumoral melatonin injection at 3% drastically inhibits tumor growth in Cal-27 xenografts after 63 days of treatment. (A, K) Tumor growth curve for mice bearing Cal-27 or SCC-9 tumors receiving vehicle (control) or intratumoral melatonin injections (0.3 [aMT 0.3%] or 3% [aMT 3%]) every day for 63 days. Cal-27 control mice were sacrificed after 41 days of treatment because tumor volume exceeded 1000 mm^3^. Data are presented as the means ± standard error of the mean (n=8-10 for each group). One-tailed unpaired *t*-test: \*\*\**P* < .001 vs. control group and ^##^*P* < .01 vs. melatonin 0.3% treated tumors. (B) Melatonin levels in tumors after 35 days of treatment. (C, L) Representative axial T2-weigthed images obtained by MRI of mice bearing Cal-27 or SCC-9 tumors treated with vehicle (control; after 41 days or 63 days, respectively) or melatonin 3% intratumorally (aMT 3%; after 63 days of treatment). White arrows indicate tumor mass. (D, M) Histological analysis of HE-stained tumors. Collagen capsule using PS staining (red). Mucin glandular structures are stained by AB (blue). Immunohistochemical identification of proliferating cells with Ki-67 antibody (brown). Scale bar = 100 µm. Asterisks mark central tumor cysts and black arrows indicate the capsule stained red by PS. (E, F, G, H, I, J, N, O, P, Q, R, S) Quantitative histochemical analyses of PS, AB, and Ki-67 staining, showing an increase in the PS collagen capsule and a decrease in AB and Ki-67 staining after 3% melatonin treatment. Experimental groups include control tumors (mice treated with vehicle) and tumors treated with 3% melatonin intratumorally for 35 days (D, E, F, G, M, N, O, P) or 63 days (D, H, I, J, M, Q, R, S). Data are presented as the means ± standard error of the mean (n=3 for each group). One-tailed unpaired *t*-test: \**P* < .05, \*\**P* < .01 vs. control tumors.

### 3.3. Intratumoral melatonin treatment potentiates CDDP effects to decrease HNSCC tumor growth

Next, we examined the capacity of the intratumoral injection of melatonin to potentiate the cytotoxic effects of CDDP in Cal-27 xenografts. Melatonin 3% was administered intratumorally every 24 hours and CDDP intraperitoneally once per week for 49 days. Intratumoral melatonin potentiated the effects of CDDP on tumor growth (Figs. 3A and S1B), which is in contrast to the results obtained with subcutaneous melatonin administration (Fig. 1D). Tumors treated with intratumoral melatonin plus CDDP began to show a decrease in volume from day 15 and had a significant decrease after 35 days of treatment and for the follow-up period. After 35 days, tumors treated with CDDP alone continued to grow (Figs. 3A, B, S1B). Strikingly, there were no tumors in mice that received the combined treatment after 49 days, as shown on MRI (Fig. 3C), which highlights the great clinical potential of melatonin to potentiate the effects of CDDP.

**Fig. 3.**
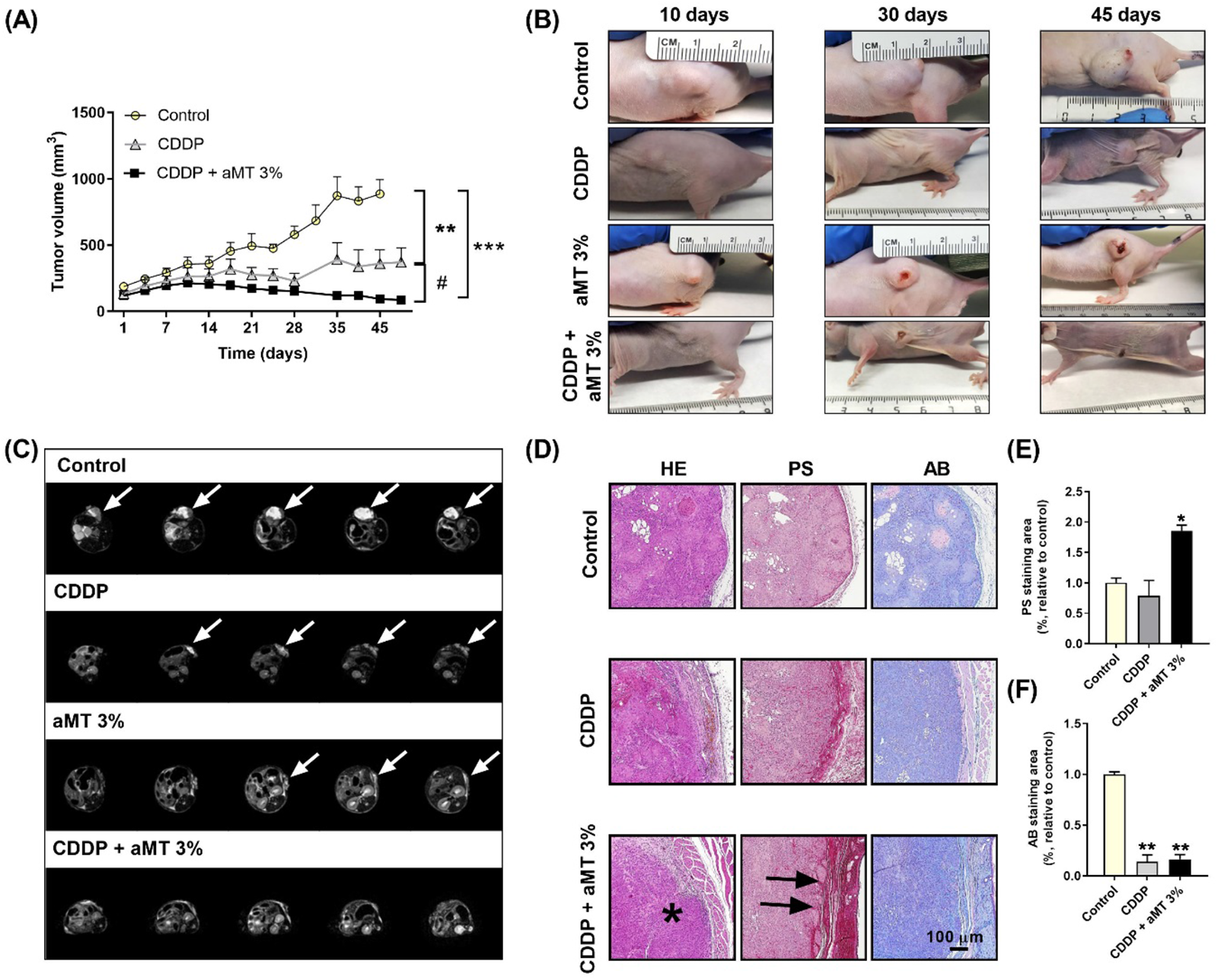
Cal-27 tumors disappear after 49 days of treatment with intratumoral melatonin and CDDP. (A) Tumor growth curve for mice bearing Cal-27 tumors treated with vehicle (control), cisplatin at 4 mg/kg day once per week (CDDP), and the combination of CDDP plus intratumoral melatonin at 3% (CDDP + aMT 3%). Data are presented as the means ± standard error of the mean (n=5-8 for each group). One-tailed unpaired *t*-test: \*\**P* < .01, \*\*\**P* < .001 vs. control group and ^#^ *P* < .05 vs. CDDP treated tumors. (B) Representative images acquired after 10, 30, and 45 days of treatment. (C) Representative axial T2-weigthed images obtained by MRI of mice bearing Cal-27 tumors treated with vehicle (control), CDDP, intratumoral injections of melatonin at 3%, and the combination of CDDP + aMT 3%. White arrows indicate tumor mass. (D) Histological analysis of HE-stained tumors. Collagen capsule using PS staining (red). Mucin glandular structures are stained by AB (blue). Scale bar = 100 µm. Asterisks mark central tumor cysts, and black arrows indicate the capsule stained red by PS. (E, F) Quantitative histochemical analyses of PS and AB staining, showing an increase in PS collagen capsule and a decrease in AB staining area after 35 days of treatment with the combination of CDDP plus melatonin at 3%. Experimental groups include control tumors (mice treated with vehicle) and tumors treated with CDDP or the combined treatment of CDDP + aMT 3% for 35 days (B, C, F). Data are presented as the means ± standard error of the mean (n=3 for each group). One-tailed unpaired *t*-test: \**P* < .05, \*\**P* < .01 vs. control tumors.

In line with the above results, histology of co-treatment with intratumoral melatonin and CDDP revealed a reduction in tumor development, with a significant increase in the collagen-rich capsule (Fig. 3D, E). CDDP, as well as the melatonin plus CDDP group, resulted in a significant reduction in the adenosquamous areas (Fig. 3D, F).

**Fig. 4.**
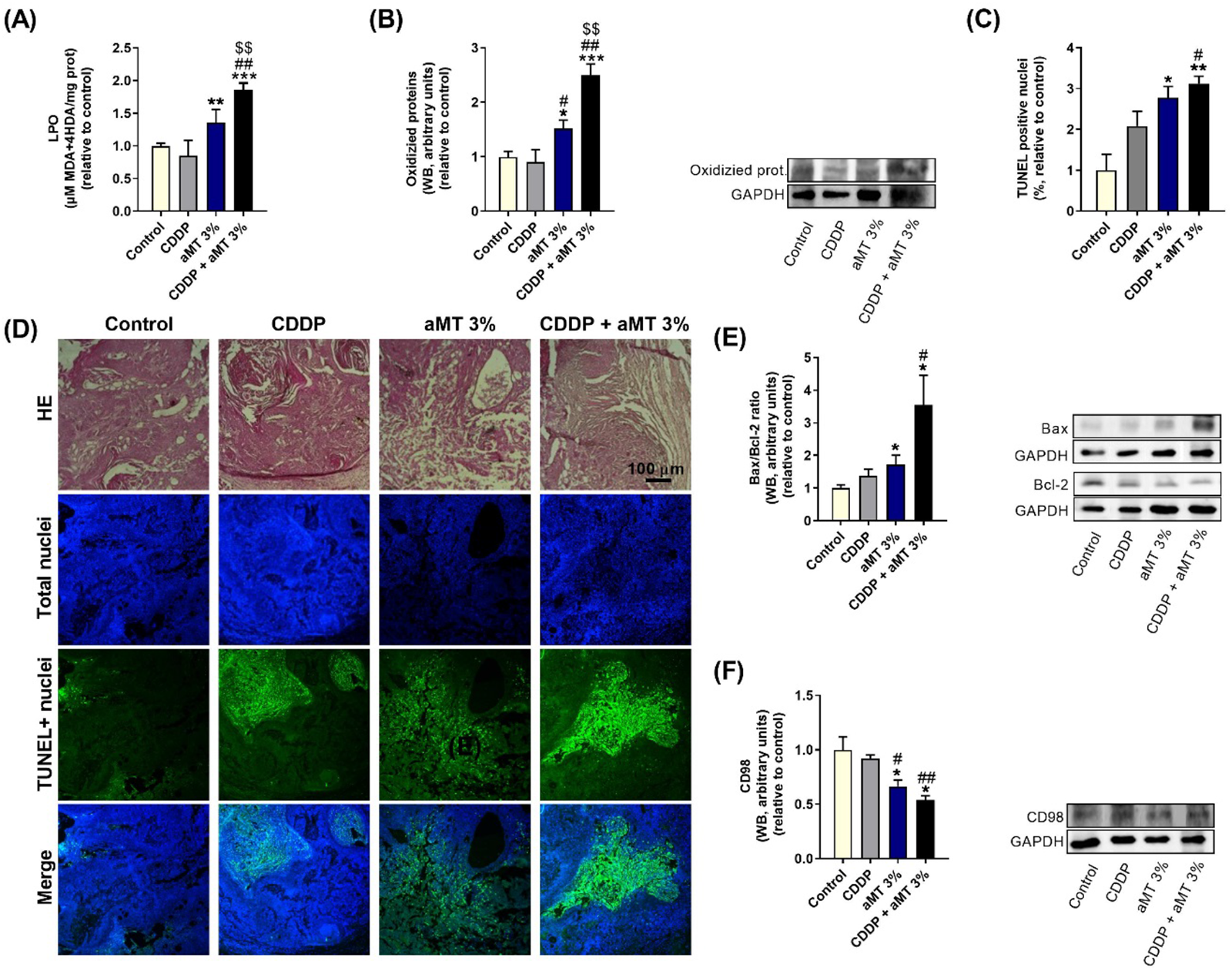
Intratumoral melatonin at 3% increases oxidative stress and apoptosis in Cal-27 xenografts tumors. (A, B) Study of oxidative stress levels and (C-E) apoptosis in Cal-27 tumors after 35 days of treatment. (A) LPO evaluation by spectrophotometric techniques. (B) Western blot (WB) analysis of oxidized proteins. (C) Percentage of apoptotic cells designated as the apoptotic index and analyzed by TUNEL assay. (D) TUNEL+ nuclei (apoptotic nuclei, green), DAPI stained nuclei (total nuclei, blue), and HE staining of the corresponding zone of the analyzed tumor. Scale bar = 100 µm. (E) Analysis of Bax/Bcl-2 expression by WB. (F) CD98 expression measured by WB. Experimental groups include control (mice treated with vehicle) and tumors from mice treated with CDDP at 4 mg/kg day once per week (CDDP), melatonin at 3% intratumorally (aMT 3%), or with the combined treatment of CDDP + aMT 3% for 35 days. Data are presented as the means ± standard error of the mean (n=4-9 for each group). One-tailed unpaired *t*-test: \**P* < .05, \*\**P* < .01, \*\*\**P* < .001 vs. control tumors; ^#^ *P* < .05, ^##^ *P* < .01 vs. CDDP and ^$$^ *P* < .01 vs. aMT 3%.

These data indicate that intratumoral administration of melatonin potentiates the cytotoxic effect of CDDP, especially to decrease tumor growth and increase the collagen capsule, indicating that it could be a proper clinical approach to improve the effectiveness of existing anti-cancer therapies.

### 3.4. Melatonin stimulates ROS production and apoptosis in Cal-27 xenografts

Given our previous study in which melatonin induced ROS complex I RET under anaerobic metabolism and led to apoptosis from excessive ROS production in HNSCC cells [11], we investigated the contribution of intracellular ROS to the potentiation of CDDP-induced cell death. For this purpose, we analyzed oxidative stress levels. Melatonin increased LPO levels and oxidized protein expression in Cal-27 tumors (Fig. 4A, B). Interestingly, treatment with melatonin plus CDDP enhanced these pro-oxidant effects and noticeably triggered ROS-dependent apoptosis, which resulted in an increase in the Bax/Bcl-2 ratio and TUNEL-positive nuclei compared to the CDDP group (Fig. 4C, D, E).

Finally, considering that an increase in ROS is related to cell differentiation [36], we analyzed CD98 expression, a cancer stem cell marker in HNSCC [37]. In agreement with the above histological results, melatonin at 3% reduced CD98 levels compared to the control and CDDP groups (Fig. 4F), suggesting an increase in tumor cell differentiation.

**Fig. 5.**
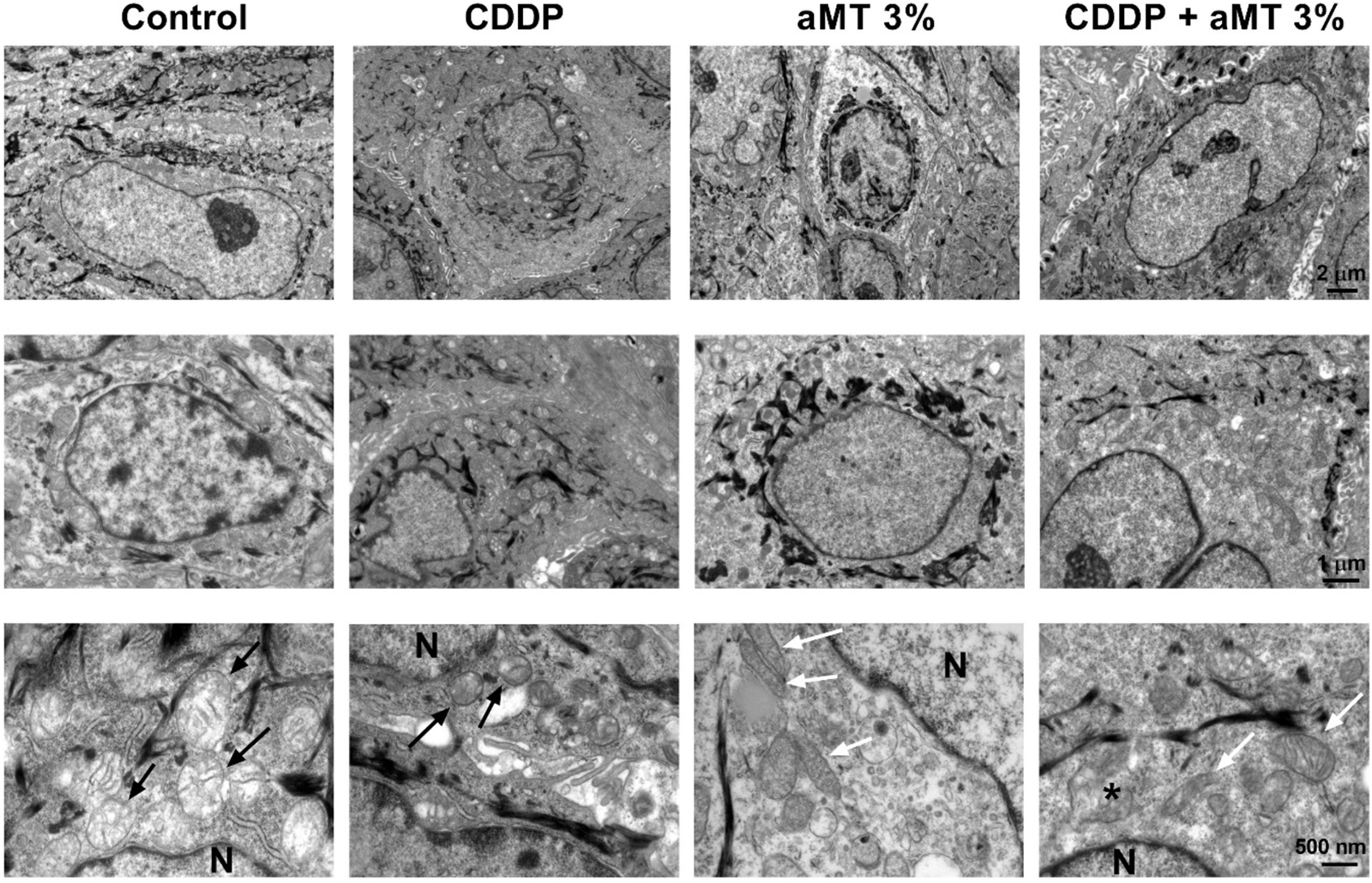
Melatonin-induced changes in mitochondrial morphology and distribution in Cal-27 xenografts tumors. Alterations in mitochondrial morphology analyzed by EM in Cal-27 xenografts. In control and CDDP tumors, the typical morphology of the Cal-27 mitochondria was observed as a round shape and few mitochondrial cristae (black arrows). Mitochondria are localized in the perinuclear region. In melatonin-treated tumors, mitochondria are elongated, with an increased number of cristae (white arrows) and following a homogenous distribution among the cellular cytoplasm. Some disruption in the mitochondrial membrane is appreciated in the melatonin-treated groups (black asterisk). Scale bar = 2 µm, 1 um and 500 nm.

### 3.5. Intratumoral melatonin promotes changes to mitochondrial morphology in HNSCC xenografts

Considering that one of the main targets of melatonin is the mitochondria, we evaluated mitochondrial morphology and distribution by electron microscopy (EM) in Cal-27 xenograft tumors. EM images showed clear changes between control and treated tumors (Fig. 5).

Mitochondria in the control and CDDP group had a spheroid and ovoid morphology with poorly developed cristae. Interestingly, in control tumors, mitochondria presented perinuclear localization (Fig. 5). The redistribution of mitochondria from the periphery of the cell to the perinuclear region has been associated with hypoxic conditions and, therefore, with tumor progression and treatment resistance [38,39].

However, intratumoral melatonin treatment resulted in changes in the mitochondrial structure, morphology, and pattern of cytoplasm localization. Mitochondria were larger with well-defined cristae compared to the control and CDDP groups (Fig. 5), which is indicative of a more developed organelle. In these tumors, mitochondria were localized in the periphery of the cells and not in the perinucleus. However, most of these elongated mitochondria had disruptions in their mitochondrial membrane, which could indicate that high doses of melatonin damaged the mitochondria (Fig. 5). These results were consistent with previous *in vitro* results [9].

**Fig. 6.**
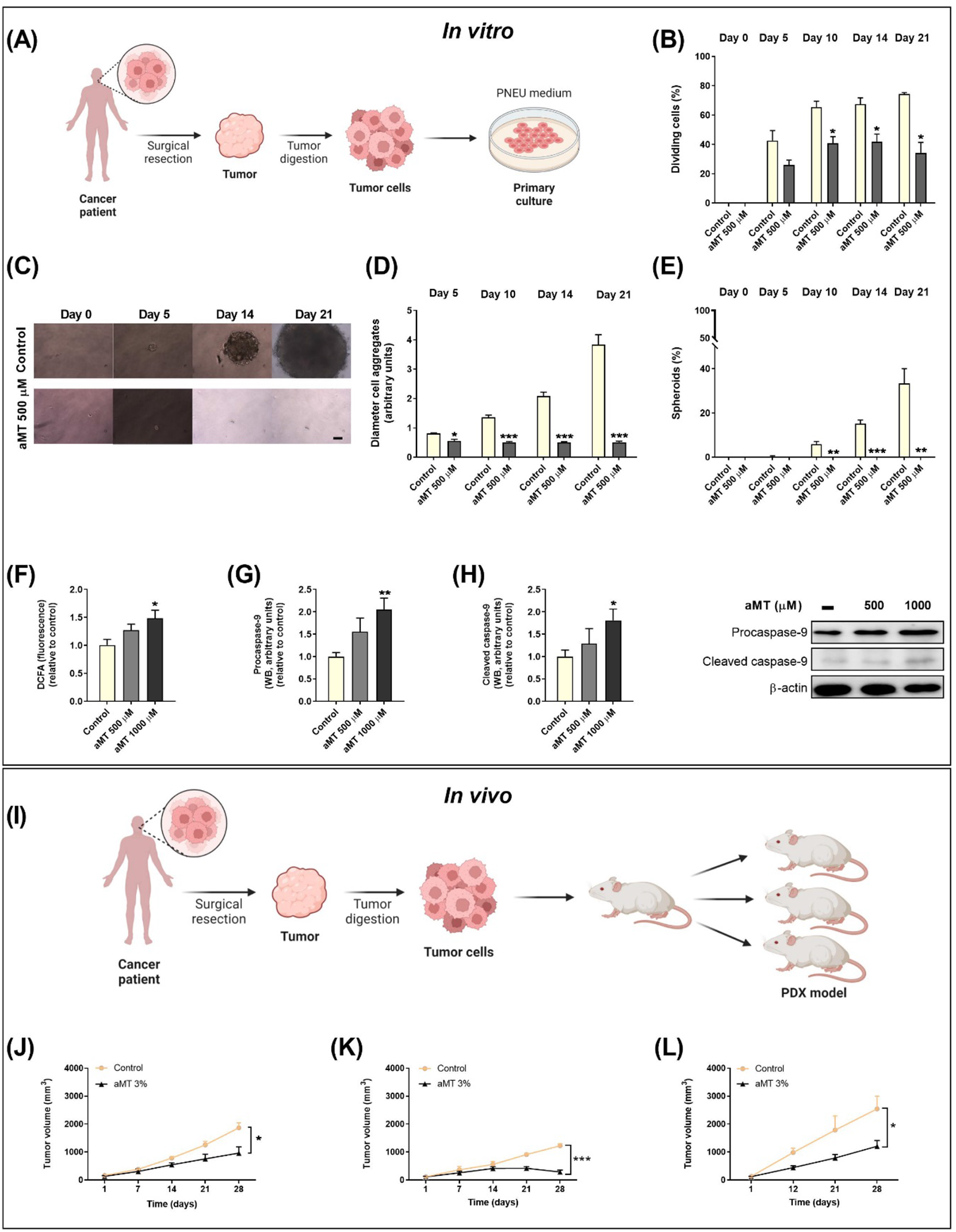
Melatonin inhibits proliferation and increases apoptosis in patient-derived tumor cells. (A) Scheme of the procedure followed to obtain a patient-derived culture (PDC). (B, C, D, E) *In vitro* proliferation analysis of SP1 cells. (B) Percentage of SP1 cells in division. (C) Representative pictures of SP1 cells. (D) Diameter of the cell aggregates formed. (E) Percentage of spheroids generated in control cells (treated with vehicle) and cells treated with melatonin at 500 µM (aMT 500 µM) after 5, 10, 14, or 21 days of treatment. Cells were treated with vehicle or melatonin at 500 µM every 72 hours to avoid cell death induced by melatonin. Data are presented as the means ± standard error of the mean (n=3 for each group). One-tailed unpaired *t*-test: \**P* < .05, \*\**P* < .01, \*\*\**P* < .001 vs. control of the respective day. (F, G, H) Evaluation of ROS production and apoptosis in SP1 cells. (F) Measurements of intracellular ROS levels by fluorimetry after staining with the DCFA fluorescent probe. (G, H) Procaspase and cleaved caspase-9 analysis by Western blot. Experimental groups include control cells (treated with vehicle) and cells treated with melatonin at 500 or 1000 µM for 48 hours. (I, J, K, L) *In vivo* analysis of melatonin’s oncostatic effect in three different patient-derived xenografts (PDXs), demonstrating that intratumoral administration of melatonin at 3% diminishes tumor growth in PDXs. (I) Scheme of the procedure followed for PDXs. (G, H, I) Tumor growth curves of mice bearing (G) SP1, (H) SP2, and (I) SP3 xenografts treated with vehicle solution (control) or the intratumoral administration of melatonin at 3% (aMT 3%). Data are presented as the means ± standard error of the mean (n=4 for each group). One-tailed unpaired *t*-test: \**P* < .05; \*\**P* < .01, \*\*\**P* < .001 vs. control group.

### 3.6. Intratumoral melatonin injection reduces tumor growth in patient-derived tumors

Next, we examined the oncostatic effect of melatonin in primary tumors derived from patients. The PDC obtained from a human biopsy of head and neck cancer tissue are referred to here as sample 1 (SP1) [23]. Primary HNSCC cells require specific culture medium and conditions to avoid cancer-associated fibroblast (CAF) overgrowth [20]. PNEU-medium provides a selection advantage to epithelial tumor cells and facilitates their proliferation so that they are not overgrown by CAFs [23]. Consequently, after patient-derived tumor tissue digestion, the obtained SP1 cells were grown in PNEU-medium (Fig. 6A). After serial passages, a culture free of CAFs was obtained and these SP1 cells were used for subsequent experiments. SP1 cells grow by forming cell aggregates, which culminate in spheroid formation. These spheroids are aggregates of single cells that undergo self-assembly [40]. Melatonin at 500 µM decreased the SP1 division rate, reducing the percentage of dividing cells from day 10 to day 21 (Fig. 6B), as well as the diameter of the cell aggregates formed (Fig. 6C, D). Interestingly, spheroid formation, which is a CSC proliferation marker [23,41], was completely abolished with melatonin 500 µM (Fig. 6C, E). Melatonin 1000 µM also increased ROS production and caspase-9 expression in SP1 primary culture (Fig. 6F-H).

Moreover, an *in vivo* study with PDX was performed from human biopsies of head and neck cancer tissues. After the digestion of tumor tissue, the obtained cells were directly transplanted into NSG mice. When tumors reached an appropriate volume, approximately 800 mm^3^, the mice were sacrificed and tumors extracted, digested, and transplanted again into 3-5 new mice to obtain a proper number of tumors to allow experimentation (Fig. 6I).

Three different tumors derived from human biopsies were used in this study: SP1, SP2 and SP3. As shown in Figure 6, the intratumoral administration of melatonin 3% reduced tumor growth compared to the control group in the three primary tumors evaluated (Fig. 6J-L). Melatonin treatment was especially effective in SP2 primary tumors because, in this PDX model, melatonin not only exhibited oncostatic effects, but was also antitumoral. SP2 tumors practically disappeared after 28 days of melatonin treatment (Fig. 6K), which is more effective than the 63 days in established Cal-27 xenografts.

### 3.7. Melatonin reduces cell migration in vitro

As high levels of ROS enable metastatic progression [42], the effect of melatonin on cell migration was explored in Cal-27 and SCC-9 cells (Fig. 7). Melatonin significantly reduced the wound healing area in a dose- and time-dependent manner (Fig. 7A-D). These results were corroborated with a Transwell system in which we observed that melatonin impaired the invasive capacities of both Cal-27 and SCC-9 cells, reducing the number of migrated cells after 48 hours of melatonin treatment (Fig. 7E-H).

Moreover, the expression of two epithelial-to-mesenchymal transition markers, vimentin and E-cadherin, which represent a better model for evaluating metastasis [43], was evaluated in SP1 primary cells. Melatonin drastically inhibited vimentin levels in SP1 cells at 500 and 1000 µM but increased E-cadherin at the highest dose (Fig. 7I, J). Overall, these data demonstrate melatonin’s ability to reduce tumor cell migration and invasion in both established and primary cells.

**Fig. 7.**
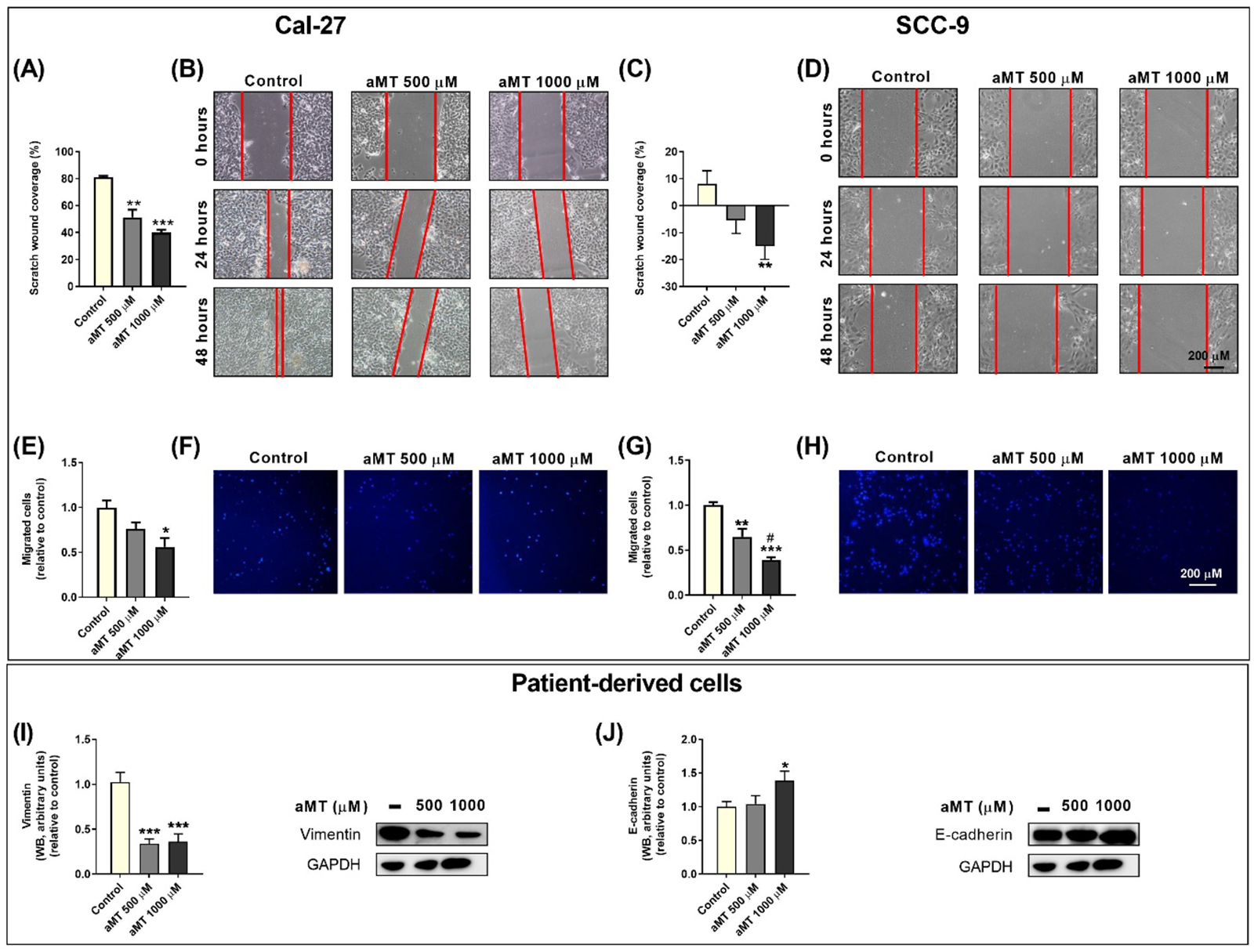
Melatonin impairs migration and invasive capacities in Cal-27 and SCC-9 cells and inhibits metastasis markers in Bi-18 cells. (A-D) Cell migration analysis of Cal-27 (A, B) and SCC-9 (C, D) cells by wound healing assay. (A, C) Quantitative analysis of scratch wound coverage and (B, D) representative images of scratched and recovering wound areas (marked by red lines) on confluence monolayers of cancer cells 0, 24, and 48 hours after wound generation. (E-H) Study of invasive capacities of Cal-27 (E, F) and SCC-9 (G, H) cells using the Transwell system. (E, G) Quantitative analysis and (F, H) representative images of migrated cells after 48 hours of melatonin treatment. Experimental groups include control (cells treated with vehicle) and cells treated with melatonin at 500 or 1000 µM (aMT 500 and 1000 µM) for 48 hours. Data are presented as the means ± standard error of the mean (n=3-5 for each group). One-tailed unpaired *t*-test: \**P* < .05, \*\**P* < .01, \*\*\**P* < .001 vs. control; ^#^ *P* < .05 vs. aMT 500 µM.

### 3.8. Melatonin reduces chemo and radiotherapy side effects in healthy organs

Intratumoral injection of melatonin seems to be safe because intratumoral melatonin does not reach the plasma (Fig. S2). However, melatonin administered subcutaneously counteracts the cytotoxic effects of IR and CDDP in healthy tissue, decreasing LPO and NOx in the liver and kidneys, as well as NOx in the heart, which was increased by IR (Fig. S2).

In addition, mice were fed 600 mg melatonin/day for 6 months to analyze subchronic toxicity. Melatonin reduced triglyceride, acid uric, creatinine, and bilirubin levels in plasma, which were increased compared to the 3-month control group (Fig. S3A, C, D, H). The enzyme GPT was also reduced after 6 months of melatonin diet (Fig. S3F). Therefore, there is no evidence available on deleterious effects from overdosing of melatonin, and this provides an opportunity for refining dose and schedule.

## 4. Discussion

Despite numerous studies demonstrating the anticancer effects of melatonin [44,45], contradictory results have been reported when it is applied *in vivo* [12]. Different melatonin treatments have been evaluated with different, controversial results [46–48]. However, to the best of our knowledge, this is the first study proposing intratumoral administration of melatonin for the treatment of HNSCC, providing evidence that melatonin can exert its oncostatic effect only by reaching a high concentration in the tumors. We observed that melatonin administered subcutaneously did not have significantly reduce tumor growth alone or in combination with IR or CDDP. These data were not in line with other studies in which melatonin was proposed as an adjuvant therapy to chemo and radiotherapy, providing synergistic antitumoral outcomes and relieving drug resistance [49–54]. The apparent discrepancies between our results and other studies were due to the melatonin levels reaching tumors. Our data suggest that melatonin can exert an oncostatic effect only at high concentrations in tumors.

Next, given that intratumoral administration was recently developed in immunotherapy [55], we decided to evaluate intratumoral injection of melatonin to increase its bioavailability in the tumor. Intratumoral injection was extremely efficacious, thereby establishing the proof of concept for intratumoral melatonin administration. Moreover, the significant increase in the collagen capsule and reduction in the observed adenosquamous areas have been related to a better cancer outcome [33–35]. Furthermore, our data do not exclude the possibility of combining melatonin with other agents, though we selected intraperitoneal CDDP. Melatonin synergized the effects of cisplatin, stimulating apoptosis and increasing oxidative stress. The increase in CDDP cytotoxicity is partly due to enhanced mitochondrial function. Melatonin induced mitochondrial elongation and facilitated the formation of cristae in HNSCC xenografts. The cristae shape has also been demonstrated to regulate the stability and assembly of respiratory chain supercomplexes, which affects respiratory efficiency, increasing ROS levels [9]. We previously demonstrated that melatonin induces ROS complex I RET and leads to apoptosis from excessive ROS production, but only at high doses [11]. Mitochondrial elongation correlates with a clear metabolic shift from glycolysis to oxidative phosphorylation that, in turn, induces ROS-dependent mitochondrial uncoupling. Thus, our results indicate that the combined treatment could offer some advantages in terms of the clinical translation of this therapeutic approach. Therefore, all things considered, our study opens the way to other intratumoral therapy combinations that may include melatonin, even immunotherapy.

On the other hand, we must consider the genetic and epigenetic differences between cell lines and original tumors, which make it difficult to evaluate how much of the original tumor biology is retained in established cell line models that have been maintained long-term *in vitro* [19,20,56]. However, culturing cancer cells from human solid tumors has generally not been rapid or readily feasible because biopsy material is usually scant and it is difficult to eliminate CAFs [56]. Nevertheless, in this study, cells from a patient’s tumor were cultured in PNEU-medium, which provides a selection advantage to epithelial tumor cells, avoiding CAF overgrowth [23]. As expected, melatonin at high doses decreased the proliferation rate and increased ROS levels and apoptosis in patient-derived cell culture, as well as completely abolishing spheroid formation, which are considered a CSC marker [23,41]. These results appear to be congruent with other studies showing that melatonin suppresses CSC-like phenotypes in other types of cancer [57,58]. Moreover, intratumoral melatonin also reduced tumor volumes in PDXs, which represent a promising pre-clinical model that can be used to predict drug responses [18].

The complex role of ROS in mediating tumor progression remains controversial. Several studies have suggested that increasing production of mitochondrial ROS enhances the metastatic capacity of tumor cells [42]. Melatonin has also been demonstrated to induce ROS complex I RET [11] and to increase apoptosis, and loss of stemness in HNSCC [59]. This is in line with our finding that melatonin increased ROS levels and suppressed the migration and invasion of HNSCC cells *in vitro* and regulated EMT-related markers in the PDC model. Therefore, the reduction of invasive capacities of HNSCC cells exerted by melatonin could be due to a reducted CSC population within these tumors. However, more experiments are needed to understand the mechanism of action of melatonin in metastasis.

## 5. Conclusions

Despite recent progress in head and neck squamous cell carcinoma treatment, the prognosis for many patients remains devastating. Moreover, primary tumor models represent a preferred approach, as research data indicates that these cells might reflect the original tumor’s properties much closer than decade old cell lines. Our study elucidated the roles of intratumoral melatonin in reducing tumor growth and synergizing CDDP and IR, and could provide new strategies for HNSCC therapy, particularly for CDDP resistance. We propose considering intratumoral co-injections of melatonin with several agents with potential for synergy.

## Author contributions

*Conceptualization*: Germaine Escames. *Methodology*: Germaine Escames, Laura Martinez-Ruiz, Javier Florido, and César Rodriguez-Santana. *Formal analysis*: Germaine Escames, Laura Martinez-Ruiz, and Javier Florido. *Investigation*: Laura Martinez-Ruiz, Javier Florido, César Rodriguez-Santana, Alba López-Rodríguez, Ana Guerra-Librero, Beatriz I. Fernández-Gil, Patricia García-Tárraga, David Sánchez-Porras, José Fernández-Martínez, Amadeo Ten-Steve, Pilar González-García, and Iryna Rusanova. *Resources*: Germaine Escames, Holger Sudhoff and Felix Oppel. *Writing-original draft preparation*: Germaine Escames and Laura Martinez-Ruiz. *Writing-review and editing*: Germaine Escames, Laura Martinez-Ruiz, José Manuel Garcia-Verdugo, Felix Oppel, Holger Sudhoff, Víctor S. Carriel, and Darío Acuña-Castroviejo. *Supervision*: Germaine Escames, Felix Oppel, Holger Sudhoff, Darío Acuña-Castroviejo, and Víctor S. Carriel. *Project administration*: Germaine Escames. *Funding acquisition*: Germaine Escames.

## Acknowledgments

This study was funded by grants from the European Regional Development Fund (B-CTS-071-UGR18); Consejería de Economía, Innovación, Ciencia y Empleo, Junta de Andalucía (P18-RT-32222); Ministerio de Ciencia e Innovación/AEI: Agencia Estatal de Investigación/ 10.13039/501100011033/ y financiado por la Unión Europea “NextGenerationEU”/ PRTR (SAF2017-85903-P; PID2020-115112RB-I00), and from the University of Granada (Grant “UNETE,” UCE-PP2017-05), Spain. J. F. and L. M. are recipients of FPU fellowships from the Ministerio de Educación Cultura y Deporte, Spain. Some experiments on PDC were supported by Klinikum Bielefeld. Finally, we wish to thank San Francisco Edit (San Francisco) for proofreading the paper.

## Data availability statement

The data that support the findings of this study are available from the corresponding author upon reasonable request.

## References

1. Hardingham, N.; Ward, E.; Clayton, N.; Gallagher, R. Acute Swallowing Outcomes After Surgical Resection of Oral Cavity and Oropharyngeal Cancers With the Mandibular Lingual Release Approach. Otolaryngol. Neck Surg. 2022, 019459982211239, doi:10.1177/01945998221123925.

2. Luo, X.; Chen, Y.; Tang, H.; Wang, H.; Jiang, E.; Shao, Z.; Liu, K.; Zhou, X.; Shang, Z. Melatonin Inhibits EMT and PD-L1 Expression through the ERK1/2/FOSL1 Pathway and Regulates Anti-Tumor Immunity in HNSCC. Cancer Sci. 2022, 113, 2232–2245, doi:10.1111/cas.15338.

3. Yao, W.; Qian, X.; Ochsenreither, S.; Soldano, F.; Deleo, A.B.; Sudhoff, H.; Oppel, F.; Kuppig, A.; Klinghammer, K.; Kaufmann, A.M.;, et al. Disulfiram Acts as a Potent Radio-Chemo Sensitizer in Head and Neck Squamous Cell Carcinoma Cell Lines and Transplanted Xenografts. Cells 2021, 10, 1–21, doi:10.3390/cells10030517.

4. Wang, Y.; Wang, Z.; Shao, C.; Lu, G.; Xie, M.; Wang, J.; Duan, H.; Li, X.; Yu, W.; Duan, W.;, et al. Melatonin May Suppress Lung Adenocarcinoma Progression via Regulation of the Circular Noncoding RNA Hsa_circ_0017109/MiR-135b-3p/TOX3 Axis. J. Pineal Res. 2022, 73, doi:10.1111/jpi.12813.

5. 5. Sahar Baghal-Sadriforoush; Morteza Bagheri; Isa Abdi Rad; Fattah Sotoodehnejadnematalahi Melatonin Sensitizes OVCAR-3 Cells to Cisplatin through Suppression of PI3K/Akt Pathway. Cell. Mol. Biol. 2022, 68, 158–169, doi:10.14715/cmb/2022.68.4.19.

6. Guerra, J.; Devesa, J. Usefulness of Melatonin and Other Compounds as Antioxidants and Epidrugs in the Treatment of Head and Neck Cancer. Antioxidants 2022, 11, 1–18, doi:10.3390/antiox11010035.

7. Li, Y.C.; Chen, C.H.; Chang, C. Lo; Chiang, J.Y.W.; Chu, C.H.; Chen, H.H.; Yip, H.K. Melatonin and Hyperbaric Oxygen Therapies Suppress Colorectal Carcinogenesis through Pleiotropic Effects and Multifaceted Mechanisms. Int. J. Biol. Sci. 2021, 17, 3728–3744, doi:10.7150/ijbs.62280.

8. Junior, R.P.; Chuffa, L.G. de A.; Simão, V.A.; Sonehara, N.M.; Chammas, R.; Reiter, R.J.; Zuccari, D.A.P. de C. Melatonin Regulates the Daily Levels of Plasma Amino Acids, Acylcarnitines, Biogenic Amines, Sphingomyelins, and Hexoses in a Xenograft Model of Triple Negative Breast Cancer. Int. J. Mol. Sci. 2022, 23, doi:10.3390/ijms23169105.

9. Guerra-Librero, A.; Fernandez-Gil, B.I.; Florido, J.; Martinez-Ruiz, L.; Rodríguez-Santana, C.; Shen, Y.Q.; García-Verdugo, J.M.; López-Rodríguez, A.; Rusanova, I.; Quiñones-Hinojosa, A.;, et al. Melatonin Targets Metabolism in Head and Neck Cancer Cells by Regulating Mitochondrial Structure and Function. Antioxidants 2021, 10, 1– 20, doi:10.3390/antiox10040603.

10. Cucielo, M.S.; Cesário, R.C.; Silveira, H.S.; Gaiotte, L.B.; Dos Santos, S.A.A.; de Campos Zuccari, D.A.P.; Seiva, F.R.F.; Reiter, R.J.; de Almeida Chuffa, L.G. Melatonin Reverses the Warburg-Type Metabolism and Reduces Mitochondrial Membrane Potential of Ovarian Cancer Cells Independent of MT1 Receptor Activation. Molecules 2022, 27, 1–15, doi:10.3390/molecules27144350.

11. Florido, J.; Martinez-Ruiz, L.; Rodriguez-Santana, C.; López-Rodríguez, A.; Hidalgo-Gutiérrez, A.; Cottet-Rousselle, C.; Lamarche, F.; Schlattner, U.; Guerra-Librero, A.; Aranda-Martínez, P.;, et al. Melatonin Drives Apoptosis in Head and Neck Cancer by Increasing Mitochondrial ROS Generated via Reverse Electron Transport. J. Pineal Res. 2022, 1–15, doi:10.1111/jpi.12824.

12. Shen, Y.Q.; Guerra-Librero, A.; Fernandez-Gil, B.I.; Florido, J.; García-López, S.; Martinez-Ruiz, L.; Mendivil-Perez, M.; Soto-Mercado, V.; Acuña-Castroviejo, D.; Ortega-Arellano, H.;, et al. Combination of Melatonin and Rapamycin for Head and Neck Cancer Therapy: Suppression of AKT/MTOR Pathway Activation, and Activation of Mitophagy and Apoptosis via Mitochondrial Function Regulation. J. Pineal Res. 2018, 64, 1–18, doi:10.1111/jpi.12461.

13. Fernandez-Gil, B.I.; Guerra-Librero, A.; Shen, Y.Q.; Florido, J.; Martínez-Ruiz, L.; García-López, S.; Adan, C.; Rodríguez-Santana, C.; Acuña-Castroviejo, D.; Quiñones-Hinojosa, A.;, et al. Melatonin Enhances Cisplatin and Radiation Cytotoxicity in Head and Neck Squamous Cell Carcinoma by Stimulating Mitochondrial ROS Generation, Apoptosis, and Autophagy. Oxid. Med. Cell. Longev. 2019, 2019, doi:10.1155/2019/7187128.

14. Mi, L.; Kuang, H. Melatonin Regulates Cisplatin Resistance and Glucose Metabolism through Hippo Signaling in Hepatocellular Carcinoma Cells. Cancer Manag. Res. 2020, 12, 1863–1874, doi:10.2147/CMAR.S230466.

15. Wang, L.; Wang, C.; Choi, W.S. Use of Melatonin in Cancer Treatment: Where Are We? Int. J. Mol. Sci. 2022, 23, 1–18, doi:10.3390/ijms23073779.

16. Wang, Y.; Tao, B.; Li, J.; Mao, X.; He, W.; Chen, Q. Melatonin Inhibits the Progression of Oral Squamous Cell Carcinoma via Inducing MiR-25-5p Expression by Directly Targeting NEDD9. Front. Oncol. 2020, 10, doi:10.3389/fonc.2020.543591.

17. Kamb, A. What’s Wrong with Our Cancer Models? Nat. Rev. Drug Discov. 2005, 4, 161–165, doi:10.1038/nrd1635.

18. Huo, K.G.; D’Arcangelo, E.; Tsao, M.S. Patient-Derived Cell Line, Xenograft and Organoid Models in Lung Cancer Therapy. Transl. Lung Cancer Res. 2020, 9, 2214– 2232, doi:10.21037/tlcr-20-154.

19. Cromwell, E.F.; Sirenko, O.; Nikolov, E.; Hammer, M.; Brock, C.K.; Matossian, M.D.; Alzoubi, M.S.; Collins-Burow, B.M.; Burow, M.E. Multifunctional Profiling of Triple-Negative Breast Cancer Patient-Derived Tumoroids for Disease Modeling. SLAS Discov. Adv. life Sci. R D 2022, 27, 191–200, doi:10.1016/j.slasd.2022.01.006.

20. Lê, H.; Seitlinger, J.; Lindner, V.; Olland, A.; Falcoz, P.-E.; Benkirane-Jessel, N.; Quéméneur, E. Patient-Derived Lung Tumoroids—An Emerging Technology in Drug Development and Precision Medicine. Biomedicines 2022, 10, 1677, doi:10.3390/biomedicines10071677.

21. Hasan, M.; Marzouk, M.A.; Adhikari, S.; Wright, T.D.; Miller, B.P.; Matossian, M.D.; Elliott, S.; Wright, M.; Alzoubi, M.; Collins-Burow, B.M.;, et al. Pharmacological, Mechanistic, and Pharmacokinetic Assessment of Novel Melatonin-Tamoxifen Drug Conjugates as Breast Cancer Drugs. Mol. Pharmacol. 2019, 96, 272–296, doi:10.1124/mol.119.116202.

22. Yang, C.-Y.; Lin, C.-K.; Tsao, C.-H.; Hsieh, C.-C.; Lin, G.-J.; Ma, K.-H.; Shieh, Y.-S.; Sytwu, H.-K.; Chen, Y.-W. Melatonin Exerts Anti-Oral Cancer Effect via Suppressing LSD1 in Patient-Derived Tumor Xenograft Models. Oncotarget 2017, 8, 33756–33769, doi:10.18632/oncotarget.16808.

23. Oppel, F.; Shao, S.; Schürmann, M.; Goon, P.; Albers, A.E.; Sudhoff, H. An Effective Primary Head and Neck Squamous Cell Carcinoma In Vitro Model. Cells 2019, 8, 555, doi:10.3390/cells8060555.

24. García-García, Ó.D.; El Soury, M.; González-Quevedo, D.; Sánchez-Porras, D.; Chato-Astrain, J.; Campos, F.; Carriel, V. Histological, Biomechanical, and Biological Properties of Genipin-Crosslinked Decellularized Peripheral Nerves. Int. J. Mol. Sci. 2021, 22, 674, doi:10.3390/ijms22020674.

25. Carriel, V.; Scionti, G.; Campos, F.; Roda, O.; Castro, B.; Cornelissen, M.; Garzón, I.; Alaminos, M. In Vitro Characterization of a Nanostructured Fibrin Agarose Bio-Artificial Nerve Substitute. J. Tissue Eng. Regen. Med. 2017, 11, 1412–1426, doi:10.1002/term.2039.

26. Chahbouni, M.; Escames, G.; Venegas, C.; Sevilla, B.; García, J.A.; López, L.C.; Muñoz-Hoyos, A.; Molina-Carballo, A.; Acuña-Castroviejo, D. Melatonin Treatment Normalizes Plasma Pro-inflammatory Cytokines and Nitrosative/Oxidative Stress in Patients Suffering from Duchenne Muscular Dystrophy. J. Pineal Res. 2010, 48, 282– 289, doi:10.1111/j.1600-079X.2010.00752.x.

27. Ulloa-Navas, M.J.; Pérez-Borredá, P.; Morales-Gallel, R.; Saurí-Tamarit, A.; García-Tárraga, P.; Gutiérrez-Martín, A.J.; Herranz-Pérez, V.; García-Verdugo, J.M. Ultrastructural Characterization of Human Oligodendrocytes and Their Progenitor Cells by Pre-Embedding Immunogold. Front. Neuroanat. 2021, 15, doi:10.3389/fnana.2021.696376.

28. Cebrián-Silla, A.; Alfaro-Cervelló, C.; Herranz-Pérez, V.; Kaneko, N.; Park, D.H.; Sawamoto, K.; Alvarez-Buylla, A.; Lim, D.A.; García-Verdugo, J.M. Unique Organization of the Nuclear Envelope in the Post-Natal Quiescent Neural Stem Cells. Stem Cell Reports 2017, 9, 203–216, doi:10.1016/j.stemcr.2017.05.024.

29. Bryan, N.S.; Grisham, M.B. Methods to Detect Nitric Oxide and Its Metabolites in Biological Samples. Free Radic. Biol. Med. 2007, 43, 645–657, doi:10.1016/j.freeradbiomed.2007.04.026.

30. Thompson, R.F.; Maity, A. Radiotherapy and the Tumor Microenvironment: Mutual Influence and Clinical Implications. In; 2014; pp. 147–165.

31. Zhang, X.; Zhang, H.; Zhang, J.; Yang, M.; Zhu, M.; Yin, Y.; Fan, X.; Yu, F. The Paradoxical Role of Radiation-induced <scp>cGAS-STING</Scp> Signaling Network in Tumor Immunity. Immunology 2022, doi:10.1111/imm.13592.

32. Zou, J.; Yuan, J.; Chen, H.; Zhou, X.; Xue, T.; Chen, R.; Zhang, L.; Ren, Z. Development of a Prognostic Score for Recommended Transarterial Chemoembolization Candidates with Spontaneous Rupture of Hepatocellular Carcinoma. J. Gastrointest. Oncol. 2022, 13, 1376–1383, doi:10.21037/jgo-22-531.

33. Ghirardi, V.; De Felice, F.; Rosati, A.; Ergasti, R.; Gueli Alletti, S.; Mascilini, F.; Scambia, G.; Fagotti, A. A Laparoscopic Adjusted Model Able to Predict the Risk of Intraoperative Capsule Rupture in Early-Stage Ovarian Cancer: Laparoscopic Ovarian Cancer Spillage Score (LOChneSS Study). J. Minim. Invasive Gynecol. 2022, 29, 961–967, doi:10.1016/j.jmig.2022.04.014.

34. Zhao, G.; Wang, C.; Tang, Y.; Liu, X.; Liu, Z.; Li, G.; Mei, Y. Glandular Differentiation in PT1 Urothelial Carcinoma of Bladder Predicts Poor Prognosis. Sci. Rep. 2019, 9, 5323, doi:10.1038/s41598-019-41844-4.

35. Xu, H.; Xie, L.; Liu, X.; Zhang, Y.; Shen, Z.; Chen, T.; Qiu, X.; Sha, N.; Xing, C.; Wu, Z.;, et al. Impact of Squamous and/or Glandular Differentiation on Recurrence and Progression Following Transurethral Resection for Non-Muscle Invasive Urothelial Carcinoma of Bladder. Oncol. Lett. 2017, 14, 3522–3528, doi:10.3892/ol.2017.6581.

36. Mendivil-Perez, M.; Soto-Mercado, V.; Guerra-Librero, A.; Fernandez-Gil, B.I.; Florido, J.; Shen, Y.-Q.; Tejada, M.A.; Capilla-Gonzalez, V.; Rusanova, I.; Garcia-Verdugo, J.M.;, et al. Melatonin Enhances Neural Stem Cell Differentiation and Engraftment by Increasing Mitochondrial Function. J. Pineal Res. 2017, 63, e12415, doi:10.1111/jpi.12415.

37. Martens-de Kemp, S.R.; Brink, A.; Stigter-van Walsum, M.; Damen, J.M.A.; Rustenburg, F.; Wu, T.; van Wieringen, W.N.; Schuurhuis, G.J.; Braakhuis, B.J.M.; Slijper, M.;, et al. CD98 Marks a Subpopulation of Head and Neck Squamous Cell Carcinoma Cells with Stem Cell Properties. Stem Cell Res. 2013, 10, 477–488, doi:10.1016/j.scr.2013.02.004.

38. Thomas, L.W.; Staples, O.; Turmaine, M.; Ashcroft, M. CHCHD4 Regulates Intracellular Oxygenation and Perinuclear Distribution of Mitochondria. Front. Oncol. 2017, 7, doi:10.3389/fonc.2017.00071.

39. Ye, X.-Q.; Li, Q.; Wang, G.-H.; Sun, F.-F.; Huang, G.-J.; Bian, X.-W.; Yu, S.-C.; Qian, G.-S. Mitochondrial and Energy Metabolism-Related Properties as Novel Indicators of Lung Cancer Stem Cells. Int. J. Cancer 2011, 129, 820–831, doi:10.1002/ijc.25944.

40. Ryu, N.-E.; Lee, S.-H.; Park, H. Spheroid Culture System Methods and Applications for Mesenchymal Stem Cells. Cells 2019, 8, 1620, doi:10.3390/cells8121620.

41. Ishiguro, T.; Ohata, H.; Sato, A.; Yamawaki, K.; Enomoto, T.; Okamoto, K. Tumor-Derived Spheroids: Relevance to Cancer Stem Cells and Clinical Applications. Cancer Sci. 2017, 108, 283–289, doi:10.1111/cas.13155.

42. Chang, C.-H.; Pauklin, S. ROS and TGFβ: From Pancreatic Tumour Growth to Metastasis. J. Exp. Clin. Cancer Res. 2021, 40, 152, doi:10.1186/s13046-021-01960-4.

43. Hou, X.; Du, C.; Lu, L.; Yuan, S.; Zhan, M.; You, P.; Du, H. Opportunities and Challenges of Patient-Derived Models in Cancer Research: Patient-Derived Xenografts, Patient-Derived Organoid and Patient-Derived Cells. World J. Surg. Oncol. 2022, 20, 37, doi:10.1186/s12957-022-02510-8.

44. Sánchez-Sánchez, A.M.; Turos-Cabal, M.; Puente-Moncada, N.; Herrera, F.; Rodríguez, C.; Martín, V. Calcium Acts as a Central Player in Melatonin Antitumor Activity in Sarcoma Cells. Cell. Oncol. 2022, 45, 415–428, doi:10.1007/s13402-022-00674-9.

45. Kurhaluk, N.; Tkachenko, H. Effects of Melatonin and Metformin in Preventing Lysosome-Induced Autophagy and Oxidative Stress in Rat Models of Carcinogenesis and the Impact of High-Fat Diet. Sci. Rep. 2022, 12, 1–14, doi:10.1038/s41598-022-08778-w.

46. Schernhammer, E.S.; Giobbie-Hurder, A.; Gantman, K.; Savoie, J.; Scheib, R.; Parker, L.M.; Chen, W.Y. A Randomized Controlled Trial of Oral Melatonin Supplementation and Breast Cancer Biomarkers. Cancer Causes Control 2012, 23, 609–616, doi:10.1007/s10552-012-9927-8.

47. A., S.; N.P., J.; A., P.; P., P.; A., C.; J., J.; J., K.; P., P.; S., P. Melatonin in Patients with Cancer Receiving Chemotherapy: A Randomized, Double-Blind, Placebo-Controlled Trial. Anticancer Res. 2014, 34, 7327–7337.

48. Seely, D.; Legacy, M.; Auer, R.C.; Fazekas, A.; Delic, E.; Anstee, C.; Angka, L.; Kennedy, M.A.; Tai, L.-H.; Zhang, T.;, et al. Adjuvant Melatonin for the Prevention of Recurrence and Mortality Following Lung Cancer Resection (AMPLCaRe): A Randomized Placebo Controlled Clinical Trial. EClinicalMedicine 2021, 33, 100763, doi:10.1016/j.eclinm.2021.100763.

49. Lee, J.H.; Yoon, Y.M.; Han, Y.S.; Yun, C.W.; Lee, S.H. Melatonin Promotes Apoptosis of Oxaliplatin-Resistant Colorectal Cancer Cells through Inhibition of Cellular Prion Protein. Anticancer Res. 2018, 38, 1993–2000, doi:10.21873/anticanres.12437.

50. Sakatani, A.; Sonohara, F.; Goel, A. Melatonin-Mediated Downregulation of Thymidylate Synthase as a Novel Mechanism for Overcoming 5-Fluorouracil Associated Chemoresistance in Colorectal Cancer Cells. Carcinogenesis 2019, 40, 422–431, doi:10.1093/carcin/bgy186.

51. Liu, Z.; Sang, X.; Wang, M.; Liu, Y.; Liu, J.; Wang, X.; Liu, P.; Cheng, H. Melatonin Potentiates the Cytotoxic Effect of Neratinib in HER2+ Breast Cancer through Promoting Endocytosis and Lysosomal Degradation of HER2. Oncogene 2021, 40, 6273–6283, doi:10.1038/s41388-021-02015-w.

52. Zhang, M.; Zhang, M.; Li, R.; Zhang, R.; Zhang, Y. Melatonin Sensitizes Esophageal Cancer Cells to 5-fluorouracil via Promotion of Apoptosis by Regulating EZH2 Expression. Oncol. Rep. 2021, 45, 22, doi:10.3892/or.2021.7973.

53. Lissoni, P.; Meregalli, S.; Nosetto, L.; Barni, S.; Tancini, G.; Fossati, V.; Maestroni, G. Increased Survival Time in Brain Glioblastomas by a Radioneuroendocrine Strategy with Radiotherapy plus Melatonin Compared to Radiotherapy Alone. Oncology 1996, 53, 43–46, doi:10.1159/000227533.

54. Berk, L.; Berkey, B.; Rich, T.; Hrushesky, W.; Blask, D.; Gallagher, M.; Kudrimoti, M.; McGarry, R.C.; Suh, J.; Mehta, M. Randomized Phase II Trial of High-Dose Melatonin and Radiation Therapy for RPA Class 2 Patients With Brain Metastases (RTOG 0119). *Int*. J. Radiat. Oncol. 2007, 68, 852–857, doi:10.1016/j.ijrobp.2007.01.012.

55. Alvarez, M.; Molina, C.; De Andrea, C.E.; Fernandez-Sendin, M.; Villalba, M.; Gonzalez-Gomariz, J.; Ochoa, M.C.; Teijeira, A.; Glez-Vaz, J.; Aranda, F.;, et al. Intratumoral Co-Injection of the Poly I:C-Derivative BO-112 and a STING Agonist Synergize to Achieve Local and Distant Anti-Tumor Efficacy. J. Immunother. Cancer 2021, 9, e002953, doi:10.1136/jitc-2021-002953.

56. Kodack, D.P.; Farago, A.F.; Dastur, A.; Held, M.A.; Dardaei, L.; Friboulet, L.; von Flotow, F.; Damon, L.J.; Lee, D.; Parks, M.;, et al. Primary Patient-Derived Cancer Cells and Their Potential for Personalized Cancer Patient Care. Cell Rep. 2017, 21, 3298–3309, doi:10.1016/j.celrep.2017.11.051.

57. Phiboonchaiyanan, P.P.; Puthongking, P.; Chawjarean, V.; Harikarnpakdee, S.; Sukprasansap, M.; Chanvorachote, P.; Priprem, A.; Govitrapong, P. Melatonin and Its Derivative Disrupt Cancer Stem-like Phenotypes of Lung Cancer Cells via AKT Downregulation. Clin. Exp. Pharmacol. Physiol. 2021, 48, 1712–1723, doi:10.1111/1440-1681.13572.

58. Liu, M.-T.; J. Reiter, R. The Impact of Melatonin and Carbon Ion Irradiation on Cancer Stem Cells. Nucl. Med. Biomed. Imaging 2017, 2, doi:10.15761/NMBI.1000127.

59. Shin, Y.Y.; Seo, Y.; Oh, S.J.; Ahn, J.S.; Song, M. hye; Kang, M.J.; Oh, J.M.; Lee, D.; Kim, Y.H.; Sung, E.S.;, et al. Melatonin and Verteporfin Synergistically Suppress the Growth and Stemness of Head and Neck Squamous Cell Carcinoma through the Regulation of Mitochondrial Dynamics. J. Pineal Res. 2022, 72, 1–16, doi:10.1111/jpi.12779.

